# Identification of Biomarker Genes Based on Multi-Omics Analysis in Non-Small Cell Lung Cancer

**DOI:** 10.1101/2022.09.05.506624

**Authors:** Ji Xia, Hai-bin He, Ying Liu, Yi Wang, Kun-Xian Shu, Ming-Yue Ma

## Abstract

**Background:** Non-small cell lung cancer (NSCLC) is a complex disease with a high mortality rate and a poor prognosis, but its molecular mechanisms and effective biomarkers are still unclear. Comprehensive analysis of multiple histological data can effectively exclude random events and is helpful in improving the reliability of the findings. In this study, we used three types of omics data, RNA-seq, microRNA-seq, and DNA methylation data, from public databases to explore the potential biomarker genes of two major subtypes of NSCLC.

**Results:** Through the combined differential analysis of multi-omics, we found 873 and 1378 potential high-risk genes in LUAD and LUSC, respectively. Then, we used WGCNA and PPI analyses to identify hub-genes and *LASSO* regression to construct prognostic models, and we obtained 15 prognostic genes. We also used survival analysis, univariate COX analysis, and GEO datasets to validate prognostic genes. Finally, we found ten genes associated with NSCLC, and eight of them have been reported in previous research.

**Conclusions:** In this study, two novel biomarker genes were identified: *NES* and *ESAM*. The two genes were both gene expression down-regulation and DNA methylation up-regulation, and regulated by *miR-122* and *miR-154*. Moreover, the *NES* gene can contribute to the clinical diagnosis and prognosis of NSCLC.

## Background

Cancer is one of the major causes of human death, and both the incidence and lethality of non-small-cell lung cancer (NSCLC) are at a high level among all cancers. According to the World Health Organization, up to 10 million people have died of cancer. Lung cancer accounts for 20% of all cancer deaths, and about 85% of lung cancer patients have NSCLC (https://www.who.int/en).

However, the occurrence, development, and molecular mechanisms of NSCLC are still unclear. The effective prevention and treatment of this disease are urgent problems that need to be studied and solved (1). Emerging technologies such as targeted therapy and immunotherapy can control disease progression, improve the cure rate, or reduce the mortality rate (2). However, the overall prognosis of NSCLC is still poor due to local recurrence, the metastasis of cancer cells, and many other factors (1). Therefore, the prediction of biomarkers for early diagnosis and targeted drug development has become the top priority of NSCLC research. The latest research shows that the main biomarkers of NSCLC are *EGFR, ALK, KRAS, FGFR*, and *PIK3CA* (3). However, traditional research methods cannot predict additional biomarkers in an efficient and accurate way. Additionally, the results of single omics analyses have a certain randomness and produce false positives. With the rapid development of high-throughput technologies, the reduction of sequencing costs, and the development of diversified sequencing technologies, in addition to bioinformatics analyses based on disease gene sequences and gene expression data, methods of sequencing proteome, microRNA, and methylome have also been applied to the study of complex diseases (4-7). Through the joint analysis of multi-omics data, the accurate classification of disease and identification of disease subtypes (8-11), the comprehensive exploration of the biological mechanisms of disease occurrence (12), and the prediction of disease-related biomarkers for diagnosis and treatment (13) can be achieved.

The In the study of the potential biomarkers of NSCLC, a comprehensive analysis of multiple histological data can effectively exclude random events at the individual level and is helpful in improving the reliability of the findings. Currently, some studies have identified the biomarkers of lung cancer through association analyses of multi-omics data (14-16). Selamat et al. obtained the LUAD biomarker (*LGALS4*) through a co-analysis of DNA methylation and mRNA data (14). By integrating the genomic and gene expression data of childhood acute lymphoblastic leukemia, multiple potential target genes (*BRCA1, COL1A1, ESR1, FGFR2*, etc.) were successfully identified (17). In addition, Cui et al. identified *miR-224* as a biomarker for NSCLC by a combined analysis of microRNA, mRNA, and methylation data (18).

Recently, an increasing number of researchers have recognized that DNA methylation has a great impact on the occurrence and development of cancer (19). As an epigenetic mechanism, DNA methylation plays an important role in the regulation of gene transcription (20). Abnormal DNA methylation has been found in a variety of cancers (21). In general, hypermethylation in the promoter region of a coding gene inhibits the expression of the gene, and promoter hypermethylation has been widely shown to contribute to the silencing of tumor suppressor genes, while promoter hypomethylation can lead to the up-regulation or activation of proto-oncogenes during carcinogenesis (16). Therefore, correlation analysis between gene methylation levels and expression levels can help to identify genes that are closely related to cancer pathogenesis.

Moreover, the regulatory mechanism of competing for the endogenous RNA (ceRNA) network can reveal the regulatory effect of microRNA on mRNA (5). There are many factors affecting ceRNA activity, such as the abundance of ceRNA components, microRNA binding capacity, RNA secondary structure, etc. Changes in these factors could lead to changes in the ceRNA network in molecular regulation, and this can lead to the occurrence of human diseases, including cancer (22-24). Therefore, studying the relationship between microRNA and mRNA will provide a reliable reference for the clinical treatment of cancer.

In this study, we used TCGA, UCSC, and GEO public databases and performed an association analysis of three types of omics data (RNA-Seq, DNA methylation signal value, and microRNA-Seq) to explore the molecular mechanism of disease development and identify more effective biomarkers to guide the screening, diagnosis, and treatment of NSCLC.

## Results

The analysis flow of this study is shown in Figure 1.

**Figure 1.**
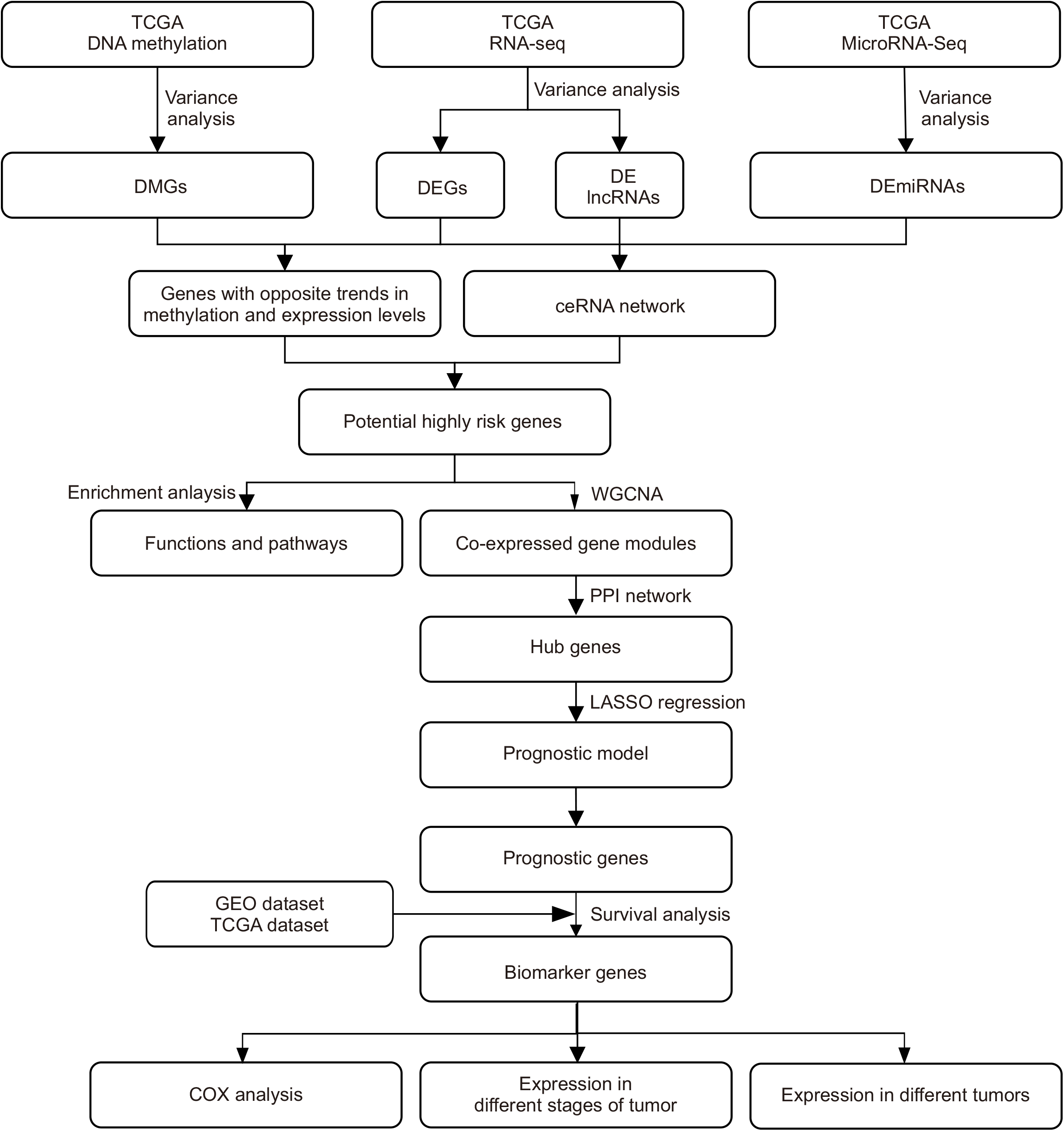
The analysis flow of this study.

### 1. Identification of Potentially High-Risk Genes by Transcriptome, RNomics, and Methylome

#### 1.1 Identification of differentially expressed genes, miRNAs, and methylations

RNA-Seq and microRNA-Seq data from the TCGA dataset were used to identify differentially expressed genes (DEGs), differentially expressed microRNA (DEmiRNAs), and differentially expressed long non-coding RNA (DElncRNAs). We obtained 5916 DEGs and 538 DEmiRNAs for LUAD, 7472 DEGs and 622 DEmiRNAs for LUSC (Table 1). DNA methylation data from UCSC Xena were used to identify differentially methylated genes (DMGs) (please see *Materials and Methods*). For LUAD, a total of 32,293 up-regulated methylation probes were distributed among 4446 DMGs, and 24,205 down-regulated methylation probes were distributed among 4783 DMGs. For LUSC, a total of 38,838 up-regulated probes were distributed among 2877 DMGs, and 63,575 down-regulated probes were distributed among 7754 DMGs. We also realized that the frequencies of up-regulated genes and miRNAs were much higher than those of down-regulated genes. Conversely, in the case of methylation genes, the frequencies of down-regulated DMGs were higher than those of up-regulated DMGs.

**Table 1.** The numbers of differentially expressed genes, microRNA, and LncRNAs in NSCLC.

| Type | DEGs |  |  | DEmiRNAs |  |  | DElncRNAs |  |  |
| --- | --- | --- | --- | --- | --- | --- | --- | --- | --- |
|  | Up | Down | Total | Up | Down | Total | Up | Down | Total |
| LUAD | 4061 | 1855 | 5916 | 418 | 120 | 538 | 1361 | 387 | 1748 |
| LUSC | 4524 | 2948 | 7472 | 472 | 150 | 622 | 1525 | 561 | 2086 |

#### 1.2 Co-expression analysis of gene expression and DNA methylation

DNA methylation levels are related to gene expression and play an important role in the occurrence and development of cancer (16). By comparing the changes in expression levels and methylation levels between cancer samples and normal samples, we found that 2656 genes and 3680 genes were related to both DEGs and DMGs in LUAD and LUSC, respectively (Supplementary Table S1).

Generally, promoters with higher methylation levels contribute to the silencing of tumor suppressor genes. Conversely, lower methylation levels lead to the activation or up-regulation of proto-oncogenes during carcinogenesis. Therefore, we identified the genes with opposite trends between gene expression levels and DNA methylation levels. Finally, 1493 genes and 2183 genes (Figure 2A and 2B; the red dots) with opposite expression differences were found. These genes were denoted as Dataset 1.

**Figure 2.**
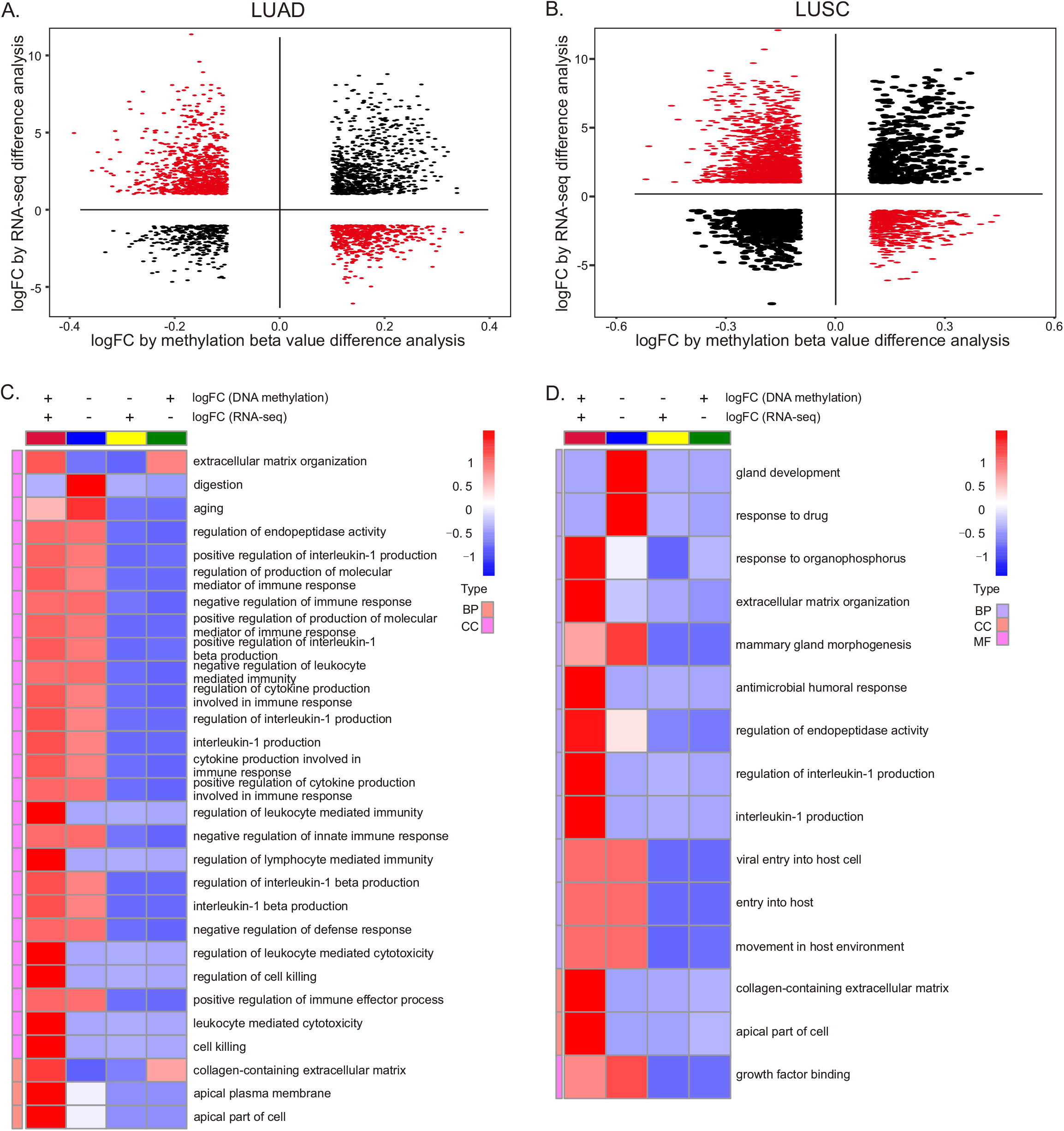
Co-expressed genes of RNA-Seq dataset and DNA methylation dataset. (A,B) Distribution of differentially expressed genes in mRNA expression levels and DNA methylation levels. The red dots represent the genes with the opposite trends, denoted as Dataset 1. (C,D) Gene enrichment analysis of co-expression genes. For the GeneIDs please see Supplementary Table S2.

Then, we compared the functions of co-expression genes. In LUAD, the genes with opposite trends prefer to enrich in immune-response-related processes, cell-killing regulation, aging, digestion, etc. (Figure 2C). In LUSC, genes with the opposite trends are likely to enrich in response to the drugs, organophosphorus, growth factor binding, gland development, etc. (Figure 2D).

#### 1.3 Construction of ceRNA regulatory network

The regulatory relationships among DEmiRNAs, DEGs, and DElncRNAs were predicted by the Starbase database. In LUAD, 122 DEmiRNAs regulate 301 DElncRNAs and 3227 DEGs. In LUSC, 113 DEmiRNAs regulate 331 DElncRNAs and 4361 DEGs. For the regulatory relationships, please see Supplementary Table S3. The DEGs involved in regulation were denoted as Dataset 2.

#### 1.4 Prediction of potential high-risk genes

In this study, the overlapping genes of Dataset 1 and Dataset 2 were defined as potential high-risk genes. Finally, a total of 878 and 1378 potential high-risk genes were obtained for LUAD and LUSC, respectively (Figure 3A and 3B).

**Figure 3.**
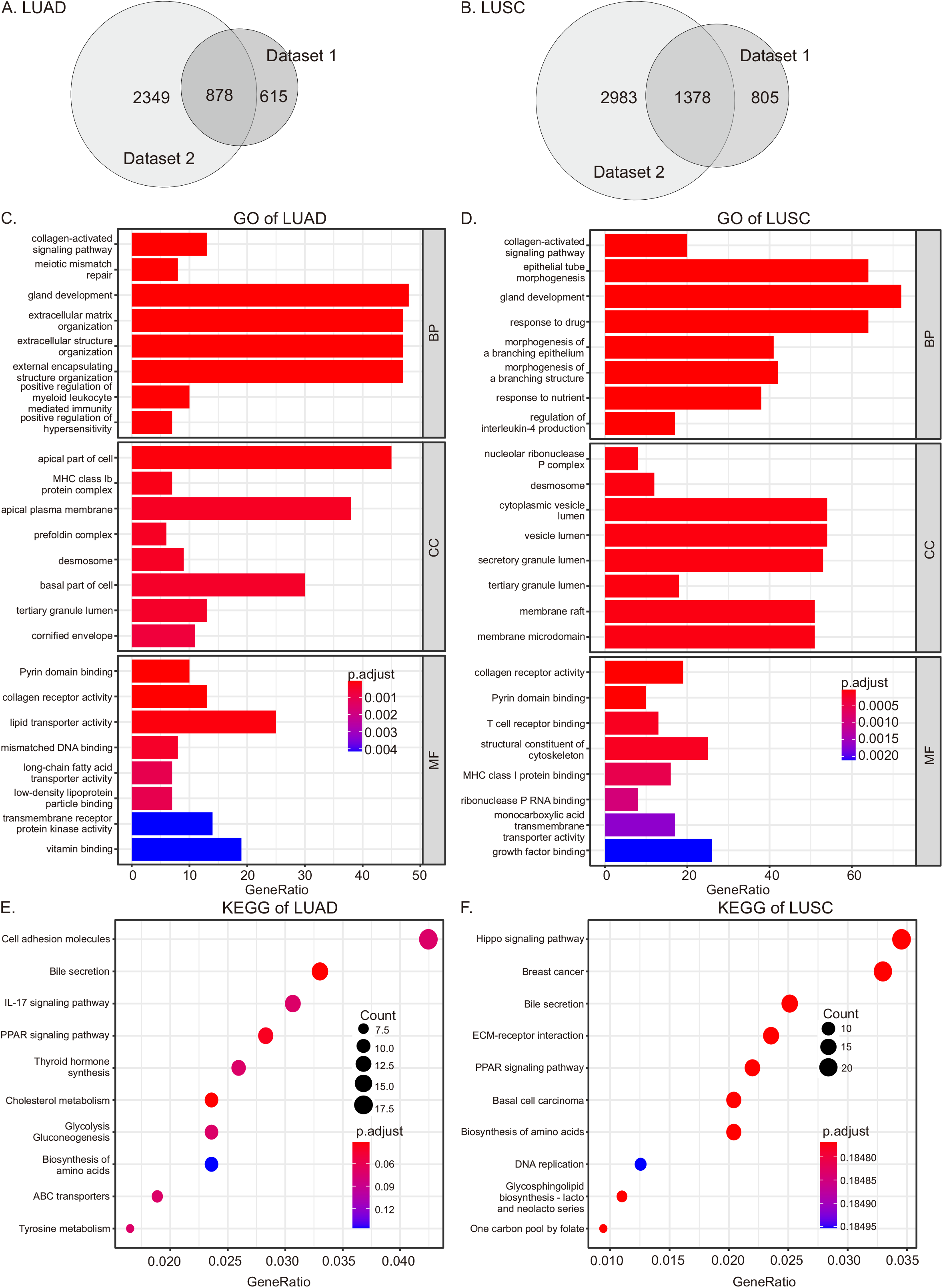
GO and KEGG enrichment analysis in NSCLC. (A,B) Venn diagram of Dataset 1 and Dataset 2. The intersection represents the potential high-risk genes. For the GeneID list, please see Supplementary Table S4. (C, D) The top 8 GO enrichment terms. BP: biological process, CC: cellular component, MF: molecular function. (E,F) The top 10 KEGG pathways. The full GO and KEGG results are shown in Supplementary Tables S5 and S6.

GO and KEGG pathway enrichment analysis showed that, in LUAD and LUSC, the potential high-risk genes were both enriched with the collagen-activated signaling pathway, gland development, Pyrin domain binding, collagen receptor activity, and the PPAR signaling pathway (Figure 3C–F). The risk genes of LUAD were also enriched in meiotic mismatch repair, the apical part of the cell, MHC class Ib protein complex, apical plasma membrane, lipid transporter activity, bile secretion pathways, and cholesterol metabolism pathways (Figure 3C, 3E). The risk genes of LUSC were related to epithelial tube morphogenesis, nucleolar ribonuclease P complex, desmosome, cytoplasmic vesicle lumen, T cell receptor binding, and basal cell carcinoma pathways, and bile secretion pathways (Figure 3D, 3F).

### 2. Prediction of Hub Genes

#### 2.1 Identification of co-expressed genes by weighted gene co-expression network analysis

For potential high-risk genes, weighted gene co-expression network analysis (WGCNA) was used to predict co-expression modules. In LUAD, we eliminated four outlier samples (Figure 4A) and identified 6 as the optimal soft threshold through the scatter plot (Figure 4B). The 878 potential high-risk genes were clustered into three modules (Figure 4C). Figure 4D shows the correlation coefficients between modules and two clinical traits; *P*-value < 0.05 for all three modules. The MEturquoise module with the highest absolute correlation coefficients (absolute correlation coefficient = 0.81, *P*-value = 3 × 10^−125^) was used for subsequent analysis, and the number of co-expressed genes was 462.

**Figure 4.**
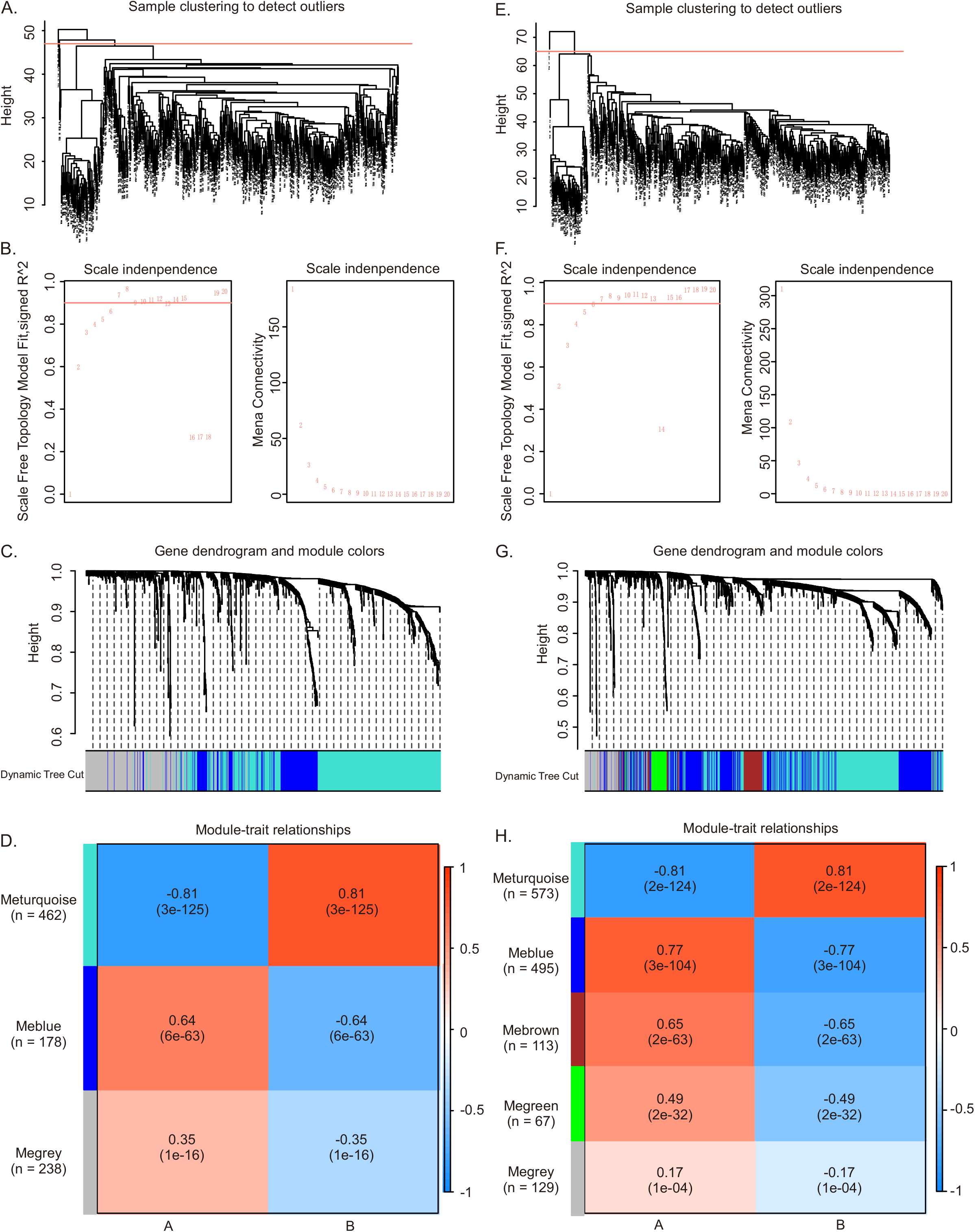
Weighted gene co-expression network analysis of potential high-risk genes in NSCLC. (a–d) for LUAD and (E–H) for LUSC. (A, E) Sample clustering. The samples with outliers were removed. (B, F) The scale-free fit index for soft-thresholding powers. The soft-thresholding power in the WGCNA was determined based on a scale-free *R*^2^ (*R*^2^ = 0.95). The left panel presents the relationship between the soft threshold and scale-free *R*^2^. The right panel presents the relationship between the soft threshold and mean connectivity. We identified 6 as the optimal soft threshold through the scatter plot. (C, G) The potential high-risk genes were clustered into different modules, presented in different colors. (D, H) The associations of each module-trait. The rows represent module eigengene, and the columns represent traits. The corresponding correlations and *P*-values were calculated by a *t*–test. For the GeneIDs of each modular gene, please see Supplementary Table S7.

In LUSC, one outlier sample was removed (Figure 4E), the optimal soft threshold was also set as 6 (Figure 4F), and the 1378 potential high-risk genes were clustered into five modules (Figure 4G).

The MEturquoise module (absolute correlation coefficient = 0.81, *P*-value = 2 × 10^−124^) was used for subsequent analysis, and the number of co-expressed genes was 573 (Figure 4H).

#### 2.2. Identification of hub genes by interaction network analysis

The STRING database was used to obtain the protein–protein interaction relationships of co-expressed genes. In LUAD, the interaction network of 462 co-expressed genes consisted of 462 nodes and 1013 edges. In LUSC, the interaction network of 573 co-expressed genes consisted of 573 nodes and 1520 edges. We used the molecular complex detection (MCODE) plugin in Cytoscape to predict hub genes. Totals of 54 and 77 hub genes were obtained for LUAD and LUSC, respectively (Figure 5).

**Figure 5.**
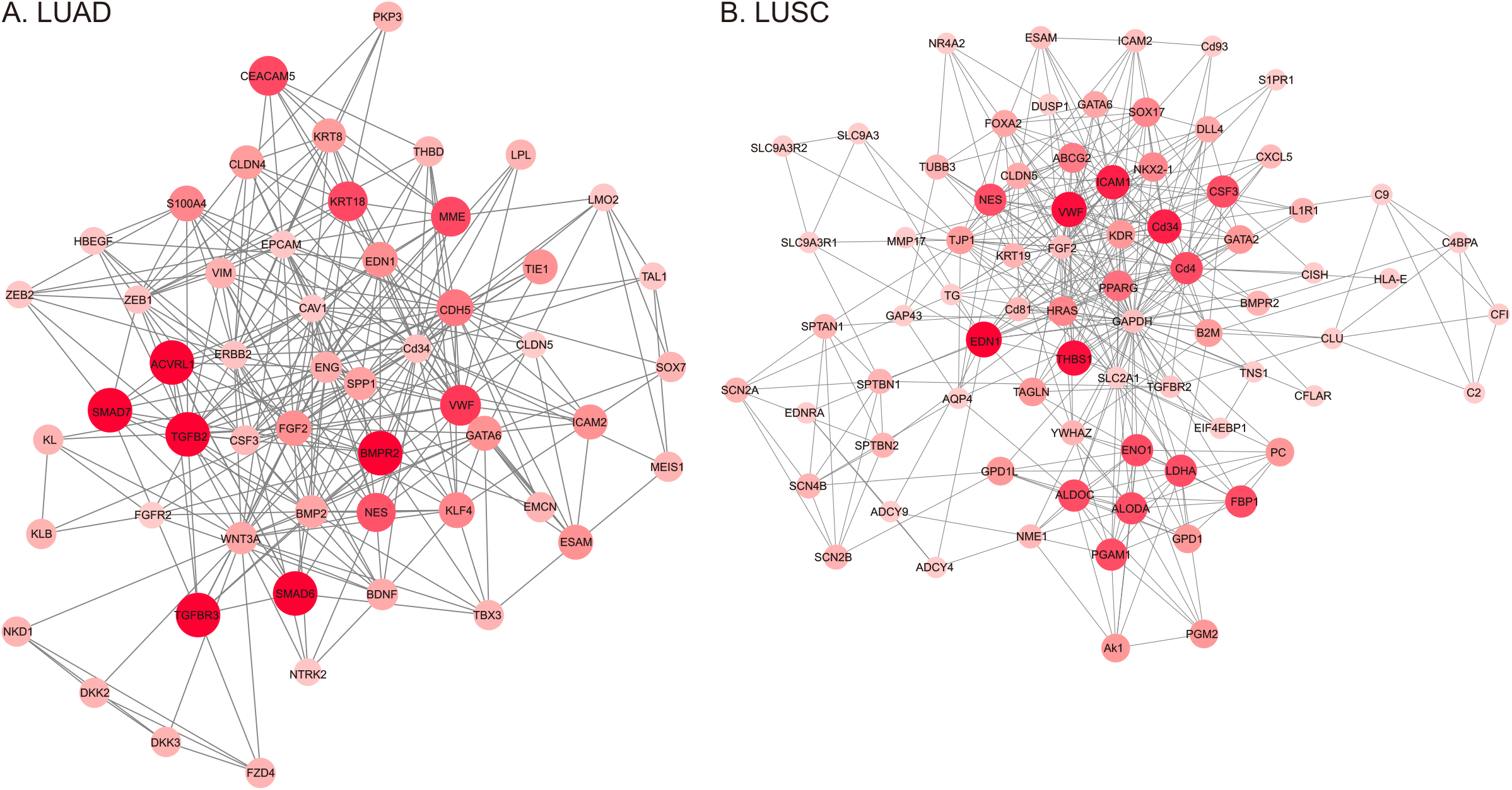
Protein interaction network of hub genes. The color and size of the nodes represent the MCODE score.

### 3. Identification of Prognostic Genes using the LASSO Regression Model

To identify the disease-characterizing genes, we used the gene expression of hub genes and the *LASSO* regression method to construct prognostic models of LUAD and LUSC. With cross-validation (nflods=20), two models were generated: the best model (*λ* = 0.0287 in LUAD, *λ* = 0.0482 in LUSC) and the simplest model (*λ* = 0.0728 in LUAD, *λ* = 0.0842 in LUSC) (Supplementary Figures S1a–d). In this study, we chose the simplest model to build the prognostic models, and 15 prognostic genes were obtained. The prognostic models were constructed based on the risk regression coefficient (Formulas 1 and 2). Then, the samples were divided into low-risk and high-risk groups based on the optimal cutoff values of each sample.

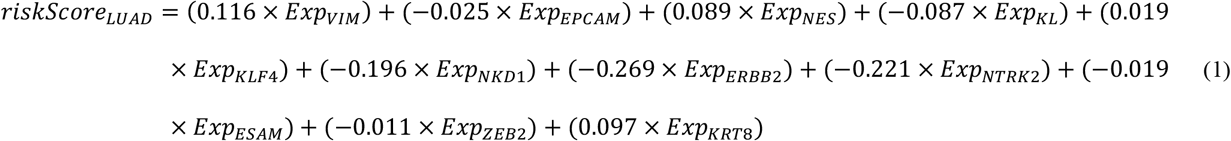

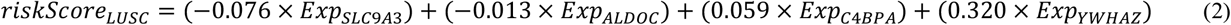

To test the accuracy of the model, we performed the following analysis on the training and testing sets, according to the above groupings. Firstly, the distribution of risk score and survival time was shown by Kaplan–Meier (KM) survival analysis and plotting the KM curve. The patients with lower risk-scores generally show better survival status and longer survival time (*Log-rank* test, *P*-value < 0.05) (Supplementary Figures S1e–h). Secondly, the distribution of risk scores for samples with different survival states (dead and alive) is shown in the boxplots. The risk scores of the patients in the dead state are significantly higher than those of the patients in the alive state (*Wilcoxon* test, *P*-value < 0.05) (Supplementary Figures S2a–d). Thirdly, the time-dependent *ROC* analysis showed that the model accuracies were 0.675 for LUAD training sets, 0.650 for LUSC training sets, 0.672 for LUAD testing sets, and 0.528 for LUSC testing sets (Supplementary Figures S2e–h).

### 4. Survival Analysis and Univariate Cox Regression Identifying Prognostic Genes Signatures

In this study, the survival analysis (*Log-rank* test) was used to test the reliability of the 15 prognostic genes as independent prognostic factors. Setting the threshold of the *P*-value to 0.05, 14 genes were related to the prognosis of LUAD and LUSC patients (Supplementary Figures S3 and S4). The survival rate of the high-expression group was higher in *EPCAM, KL, NKD1, NTRK2, ESAM, ZEB2, SLC9A3*, and *ALDOC*. Conversely, the survival rate of the low-expression group was higher in *VIM, NES, KLF4, KRT8, C4BPA*, and *YWHAZ*.

Then, a univariate COX analysis was performed on the 14 genes associated with prognosis, and the *P*-values of seven genes were less than 0.05 (Figure 6). In LUAD, *ZEB2* was identified as a protective factor, with *NES, KLF4*, and *KRT8* as risk factors. In LUSC, *ALDOC* was identified as a protective factor, with *C4BPA* and *YWHAZ* as risk factors.

**Figure6.**
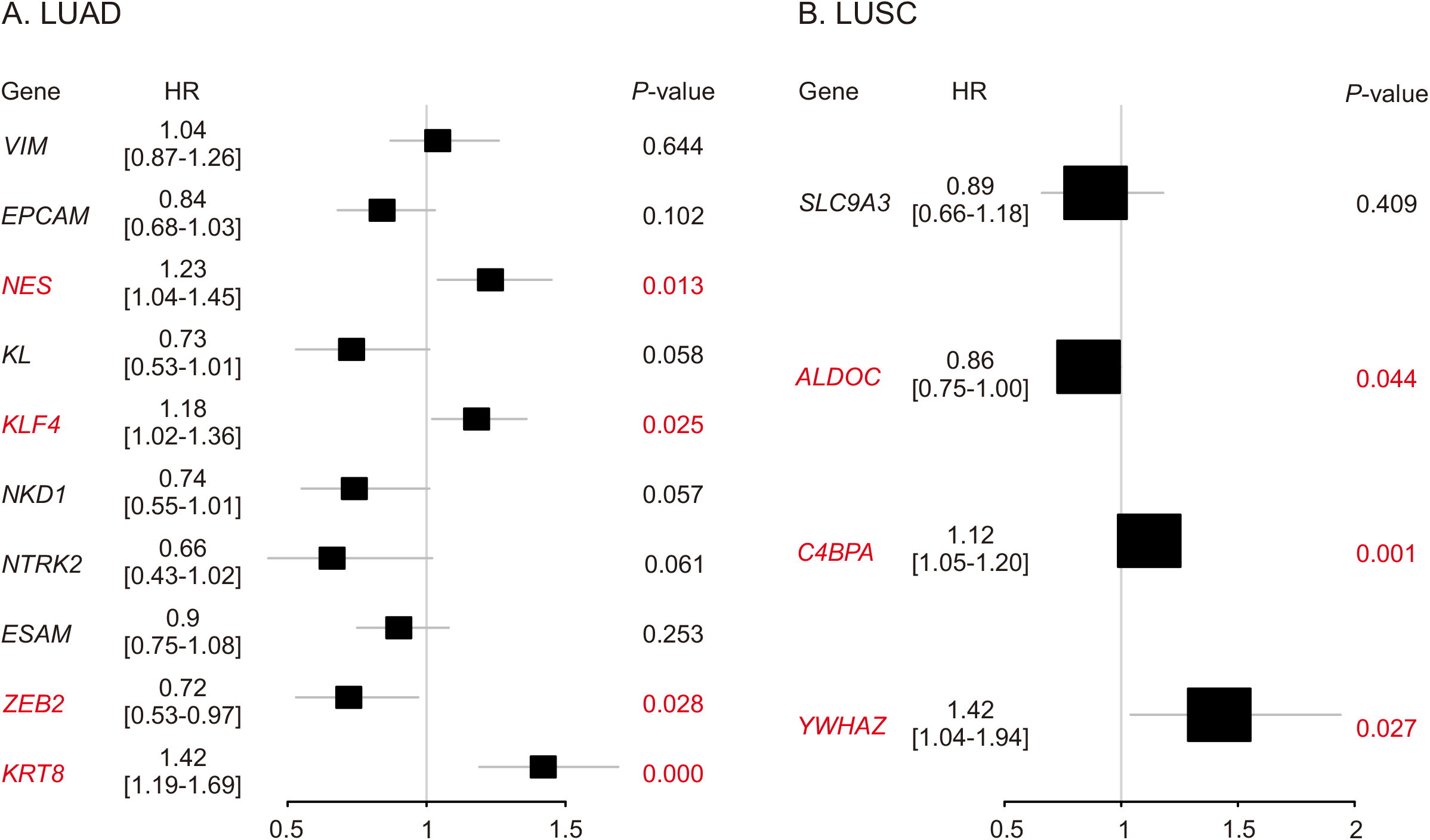
Univariate COX analysis of prognostic genes.

### 5. Validation of Prognosis-Related Genes using GEO Datasets

The 14 prognosis-related genes were again validated with GSE50081, GSE29013, and GSE37745 using an independent KM survival analysis; the threshold *P*-value was 0.05. Finally, we found ten prognosis-related genes in NSCLC (Supplementary Figure S5). Among them, the high-expression group of *NKD1, NTRK2*, and *ZEB2* had a higher survival rate. The survival rates for the other seven genes were higher in the low-expression group.

### 6. Expression of Prognostic Genes

For prognostic genes, we analyzed the gene expression level in different stages of lung cancer (Table 2) and among different tumor types (Figure 7).

**Table 2.** Intron losses and gains in Caenorhabditis

| Gene | Average expression level<sup>I</sup> (sample number) |  |  |  | P-value | P-value (Scheffe test) |  |  |  |  |  |
| --- | --- | --- | --- | --- | --- | --- | --- | --- | --- | --- | --- |
|  | Normal | Stage I | Stage II | Late Stage |  | N VS I | N VS II | N VS Late | I VS II | I VS Late | II VS Late |
| *VIM* | 7.41 (56) | 6.24 (268) | 6.20 (115) | 6.25 (101) | $4.13 \times 10^{-20}$ | $2.20 \times 10^{-18}$ | $3.75 \times 10^{-16}$ | $1.64 \times 10^{-14}$ | 0.980 | 1.000 | 0.987 |
| *EPCAM* | 6.02 (56) | 7.59 (268) | 7.65 (115) | 7.51 (101) | $7.59 \times 10^{-40}$ | $4.16 \times 10^{-37}$ | $6.28 \times 10^{-33}$ | $3.70 \times 10^{-27}$ | 0.934 | 0.839 | 0.625 |
| *NES* | 4.67 (56) | 3.05 (268) | 3.28 (115) | 3.24 (101) | $1.47 \times 10^{-26}$ | $2.01 \times 10^{-26}$ | $1.15 \times 10^{-16}$ | $1.12 \times 10^{-16}$ | 0.205 | 0.379 | 0.996 |
| *KL* | 2.21 (56) | 0.67 (268) | 0.64 (115) | 0.66 (101) | $1.77 \times 10^{-54}$ | $1.88 \times 10^{-50}$ | $2.98 \times 10^{-44}$ | $2.13 \times 10^{-41}$ | 0.961 | 0.999 | 0.991 |
| *KLF4* | 5.26 (56) | 2.98 (268) | 3.05 (115) | 3.15 (101) | $4.16 \times 10^{-39}$ | $1.17 \times 10^{-37}$ | $6.62 \times 10^{-30}$ | $1.52 \times 10^{-26}$ | 0.961 | 0.645 | 0.933 |
| *NKD1* | 1.78 (56) | 0.72 (268) | 0.65 (115) | 0.62 (101) | $2.57 \times 10^{-31}$ | $3.09 \times 10^{-27}$ | $3.55 \times 10^{-25}$ | $2.72 \times 10^{-25}$ | 0.818 | 0.627 | 0.990 |
| *ERBB2* | 4.11 (56) | 4.98 (268) | 4.93 (115) | 4.81 (101) | $1.81 \times 10^{-10}$ | $3.53 \times 10^{-10}$ | $1.39 \times 10^{-07}$ | $1.74 \times 10^{-5}$ | 0.973 | 0.446 | 0.798 |
| *NTRK2* | 1.11 (56) | 0.33 (268) | 0.29 (115) | 0.35 (101) | $2.30 \times 10^{-18}$ | $9.41 \times 10^{-17}$ | $4.09 \times 10^{-15}$ | $1.23 \times 10^{-12}$ | 0.950 | 0.994 | 0.910 |
| *ESAM* | 5.87 (56) | 3.94 (268) | 3.84 (115) | 3.83 (101) | $6.89 \times 10^{-45}$ | $4.79 \times 10^{-40}$ | $1.11 \times 10^{-36}$ | $1.51 \times 10^{-35}$ | 0.777 | 0.761 | 1.000 |
| *ZEB2* | 2.47 (56) | 1.46 (268) | 1.38 (115) | 1.35 (101) | $2.00 \times 10^{-38}$ | $5.01 \times 10^{-33}$ | $2.87 \times 10^{-31}$ | $1.63 \times 10^{-31}$ | 0.631 | 0.388 | 0.982 |
| *KRT8* | 5.97 (56) | 7.18 (268) | 7.33 (115) | 7.46 (101) | $7.32 \times 10^{-24}$ | $4.62 \times 10^{-18}$ | $1.03 \times 10^{-18}$ | $4.88 \times 10^{-21}$ | 0.443 | 0.053 | 0.780 |
| *SLC9A3* | 0.09 (48) | 0.56 (239) | 0.57 (151) | 0.54 (84) | $9.52 \times 10^{-6}$ | $2.77 \times 10^{-5}$ | $4.61 \times 10^{-5}$ | $8.17 \times 10^{-4}$ | 0.999 | 0.994 | 0.985 |
| *ALDOC* | 2.61 (48) | 4.03 (239) | 3.99 (151) | 3.89 (84) | $2.84 \times 10^{-17}$ | $1.75 \times 10^{-16}$ | $1.80 \times 10^{-14}$ | $1.28 \times 10^{-10}$ | 0.991 | 0.757 | 0.899 |
| *C4BPA* | 6.95 (48) | 2.95 (239) | 1.94 (151) | 2.74 (84) | $7.26 \times 10^{-44}$ | $4.66 \times 10^{-32}$ | $1.94 \times 10^{-43}$ | $9.87 \times 10^{-28}$ | $1.71 \times 10^{-5}$ | 0.858 | 0.028 |
| *YWHAZ* | 6.07 (48) | 7.37 (239) | 7.32 (151) | 7.44 (84) | $6.01 \times 10^{-52}$ | $2.83 \times 10^{-48}$ | $9.80 \times 10^{-42}$ | $3.80 \times 10^{-42}$ | 0.782 | 0.741 | 0.340 |

Note: We detected six putative intron gains in *C. tropicalis* and 12 in *C. sinica*. However, none of their annotations were confirmed by the RNA-seq data and thus, they were not counted as novel introns.

**Figure7.**
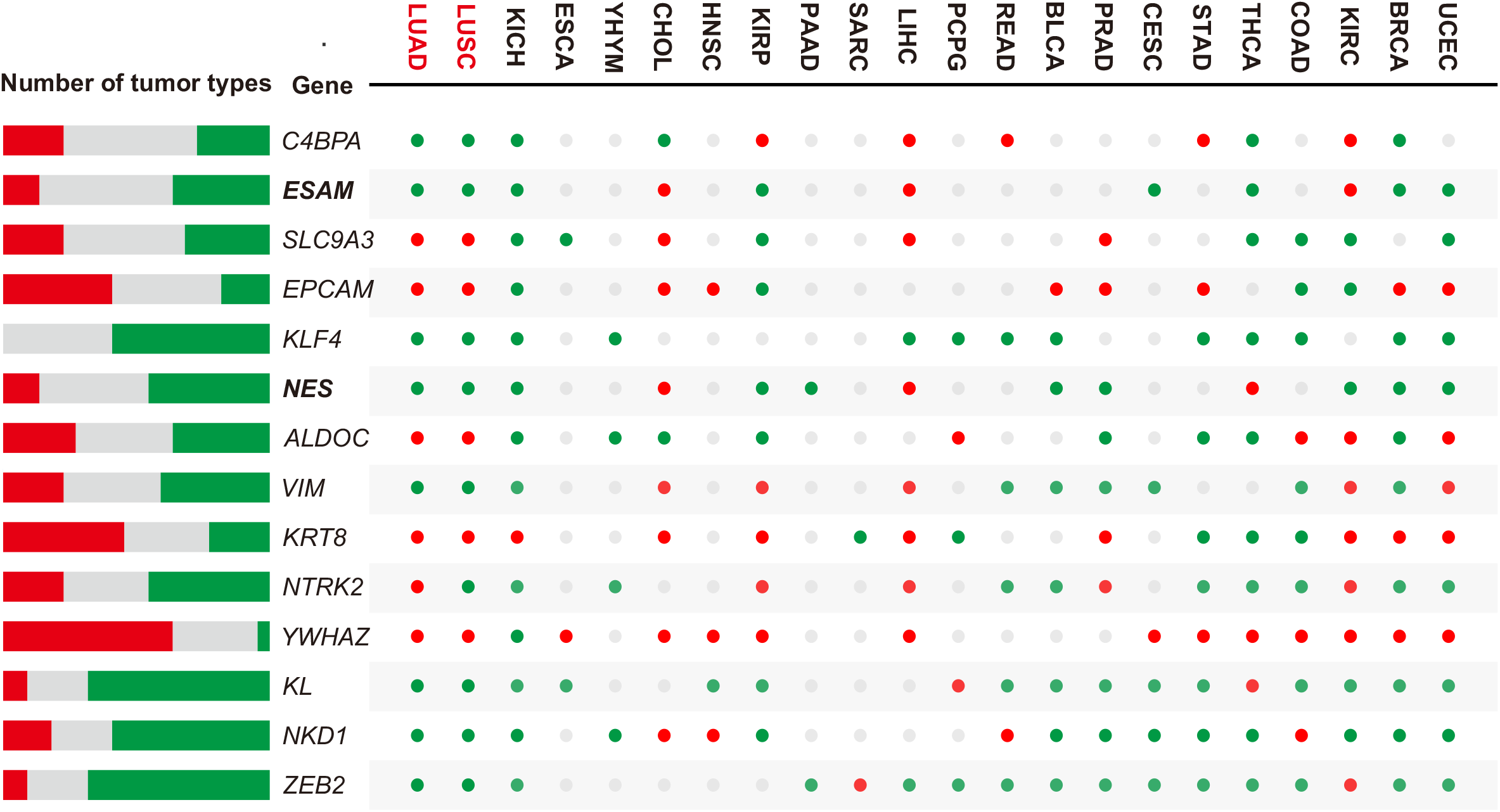
The gene expression levels in different tumor types. The red dots represent the fact that the expression of a gene in tumor samples was significantly higher than that in normal samples the green dots represent the fact that the expression of a gene in tumor samples was significantly lower than that in normal samples and the gray dots represent a lack of statistical difference.

A two-way ANOVA was used to compare the expression levels among normal samples, Stage I, Stage II, and the Late Stage. All prognostic genes showed significant differences in expression levels at different stages (Table 2, *ANOVA* test, *P*-value < 0.05 for all cases). The *Scheffe* test was used for multiple comparison tests. For the prognostic genes, there were significant differences in the expression levels between normal samples and any stage of tumor samples. However, there was almost no difference in expression levels among samples with different tumor stages. Only the expression of the *C4BPA* gene was significantly different between Stage I and Stage II, and significantly different between Stage II and the Late Stage.

After excluding the tumor types without control samples, 22 tumor types remained for comparing the differential expression of prognostic genes between tumor samples and normal samples. The expressions of different genes in different tumors were not completely consistent (Figure 7 and Figure S5). For instance, the expressions of the *KL* gene of PCPG and THCA tumor samples were significantly higher than those of normal samples (the red dots in Figure 7, *Wilcoxon* test, *P*-value < 0.05). Conversely, in LUAD, LUSC, KICH, ESCA, HNSC, KIRP, READ, BLCA, PRAD, CESC, STAD, COAD, KIRC, BRCA and UCEC, the expression levels of the *KL* gene were significantly lower than those of normal samples (the green dots in Figure 7, *Wilcoxon* test, *P*-value < 0.05).

However, there was no statistical difference in the *KL* gene expression between tumor samples and normal samples in YHYM, CHOL, PAAD, SARC, and LIHC (the gray dots in Figure 7, *Wilcoxon* test, *P*-value > 0.05).

### 7. New Biomarker Genes: NES and ESAM

In the former analysis, we identified 10 prognostic genes for NSCLC. According to previous research, *EPCAM, NKD1, NTRK2, ZEB2, VIM, KRT8, KLF4*, and *YWHAZ* have been used as prognostic markers or have been recognized as potential markers (25-32). Therefore, we found two new potential biomarker genes: *NES* and *ESAM*.

The *NES* gene is a gene expression down-regulation and DNA methylation up-regulation. A total of 28 SNP sites were all distributed in the *NES* coding region. Among of them, 61% comprised missense mutations (*N* = 17), 32% comprised silent mutations (*N* = 9), and 7% comprised nonsense mutations (*N* = 2). The univariate *COX* analysis implied that the *NES* gene was a risk factor in LUAD (Figure 6A, *hazard ratio* = 1.23, *P*-value = 0.013). The survival analyses of both the TCGA and GEO datasets revealed that the low expression of *NES* was conducive to patient survival (Figure 8A, *Log-rank* test, *P*-value = < 0.05 for all cases). By comparing the differential expression among 22 tumor types, it was found that the *NES* gene expression of tumor samples was lower than that of normal samples for LUAD, LUSC, KICH, KIRP, PAAD, BLCA, PRAD, KIRC, BRCA, and UCEC, but higher for CHOL, LIHC, and, THCA (Figure 8C).

**Figure8.**
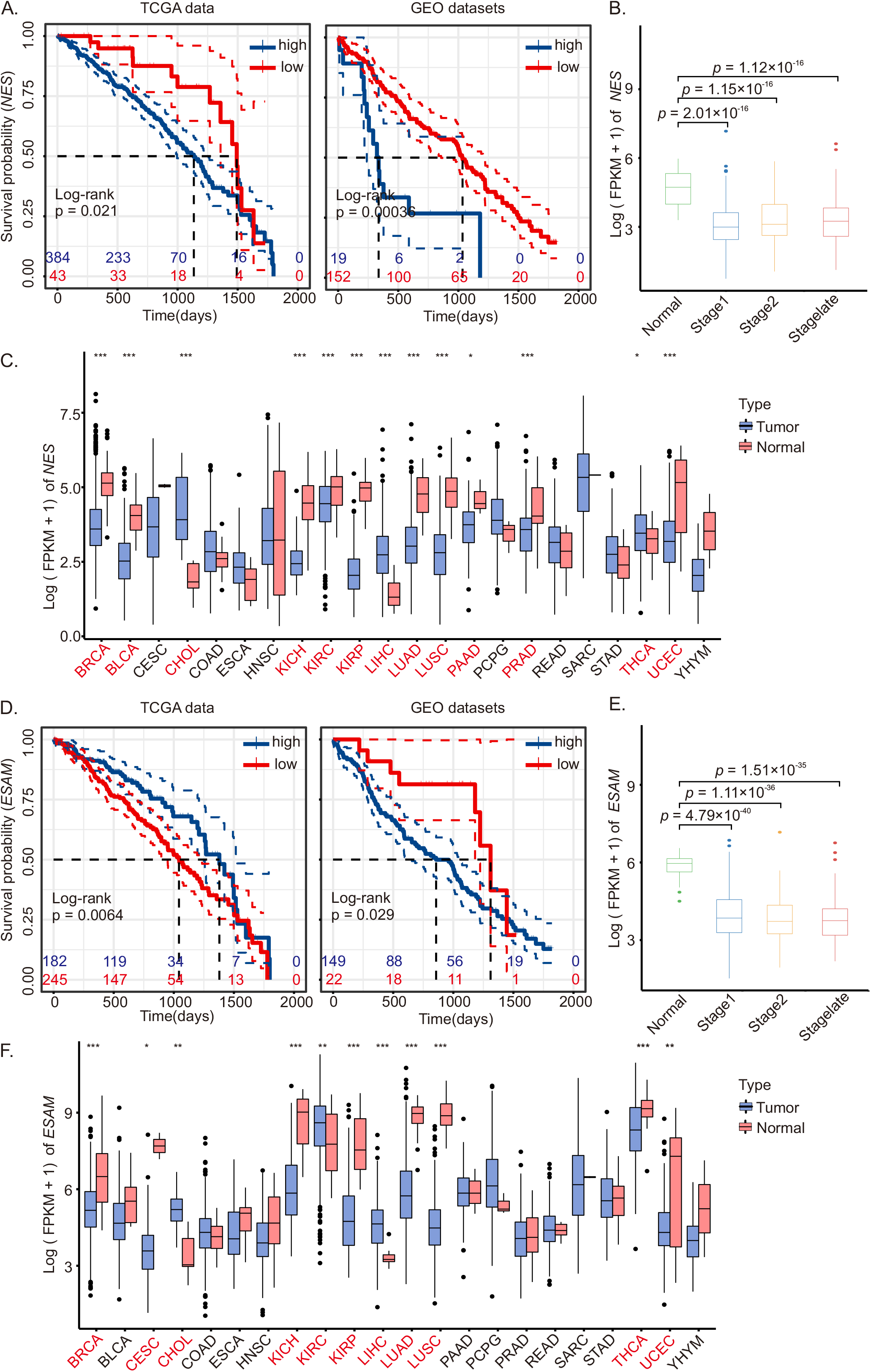
Expression analysis of individual prognostic genes. (a, d) KM curve for the TCGA dataset and GEO datasets of *NES* (a) and *ESAM* (d). (b, e) Gene expression at different tumor stages of *NES* (b) and *ESAM* (e). The *Scheffe* test was used for multiple comparison tests. (c, f) Gene expression in different types of tumors of *NES* and *ESAM*. The *Wilcoxon* test was used for statistical analysis (***: *P* < 0.001, **: *P* < 0.01, *: *P* < 0.05).

The *ESAM* gene was also a gene expression down-regulation and DNA methylation up-regulation. Only one SNP site was located in the coding region, causing a missense mutation. The *hazard ratio* value of the *ESAM* gene was 0.9, which means that *ESAM* might be one of the protective genes; unfortunately, there was no statistically significant difference (*P*-value > 0.05, Figure 6A). The results of the survival analyses based on TCGA and GEO datasets were different (Figure 8D). The TCGA data displayed the high-expression group with a good survival rate (*Log-rank* test, *P*-value = 0.006). The GEO dataset exhibited the low-expression group with a higher survival rate (*Log-rank* test, *P*-value = 0.029). Additionally, we found that *ESAM* expression was lower in the tumor samples of LUAD, LUSC, KICH, KIRP, CESE, THCA, BRCA, and UCEC, and more highly expressed in the tumor samples of CHOL, LIHC, and KIRC (Figure 8F).

## Discussion

NSCLC is a complex disease with high mortality rates and poor prognosis. However, the molecular mechanism and effective biomarkers of NSCLC are still unclear. In this study, we used three types of omics data and a variety of bioinformatics analysis methods to predict the potential biomarkers of LUAD and LUSC, which are two major subtypes of NSCLC.

Three types of omics data, transcriptome, RNomics, and methylome, were used to identify potential high-risk genes. Using co-expression analyses of RNA-Seq and DNA methylation, we defined the genes with opposite trends of the gene expression levels and DNA methylation levels as one dataset. The genes of Dataset 1 were enriched in immune-response-related processes, cell-killing regulation, aging, digestion, collagen-containing extracellular matrix, and the apical part of the cell (Figures 2c and 2d). Using RNA-Seq and microRNA-Seq data to construct the ceRNA regulatory network, we defined the genes related to regulation as another dataset. The overlapping genes of the two datasets were regarded as high-risk genes. Totals of 878 and 1378 potential high-risk genes were obtained for LUAD and LUSC, respectively. The potential high-risk genes were enriched with the collagen-activated signaling pathway, gland development, pyrin domain binding, collagen receptor activity, and the PPAR signaling pathway.

Then, bioinformatics methods and statistical analyses were used to predict the prognostic genes of NSCLC. Firstly, 462 and 573 co-expression genes of LUAD and LUSC were found by WGCNA. Secondly, using PPI analysis to investigate the interaction of the potential prognostic genes, more than 100 hub genes were obtained. Thirdly, using LASSO analysis to construct prognostic models, 15 prognostic genes were identified. The KM survival demonstrated that 14 of these genes were significantly associated with prognosis in NSCLC (*EPCAM, KL, NKD1, NTRK2, ESAM, ZEB2, VIM, NES, KLF4, KRT8, SLC9A3, ALDOC, C4BPA*, and *YWHAZ*). Next, we used a univariate COX analysis to test the independent prognostic role of the prognostic genes. Five genes (*NES, KLF4, KRT8, C4BPA*, and *YWHAZ*) were identified as risk factors, and two genes (*ZEB2* and *ALDOC*) were identified as protective factors. Lastly, we also performed a validation analysis using GEO datasets.

Finally, we identified ten genes associated with the prognosis of NSCLC.

According to former research, *EPCAM, NKD1, NTRK2, ZEB2, VIM, KRT8, KLF4*, and *YWHAZ* have been used as prognostic markers or have been recognized as potential markers for NSCLC (25-31). Therefore, we identified two new potential biomarker genes: *ESAM* and *NES*. The two genes were both gene expression down-regulation and DNA methylation up-regulation, and were simultaneously regulated by *miR-122* and *miR-154*.

The endothelial cell adhesion molecule (*ESAM*) is a member of the immunoglobulin superfamily that mediates homologous adhesion between endothelial cells. The gene is related to cell–cell adhesion mediator activity, protein binding, and other molecular functions (33, 34). In addition, some studies have shown that *ESAM*-regulated tumor metastasis through endothelial cell migration and tube formation in metastatic nodules during angiogenesis. Therefore, the inhibition of *ESAM* expression could inhibit angiogenesis and thus inhibit tumor metastasis (35). In another study, the overexpression of *miR-7* was found to inhibit the expression of *ESAM*, thereby reducing the metastasis of breast cancer stem cells (36). In this study, the differential expression of the *ESAM* gene significantly impacted the prognosis of LUAD. By comparing the expressions of the *ESAM* for different stages of cancer development, it was found that the *ESAM* expressions for any cancer period were significantly lower than those of normal samples. However, we found that the high expression of *ESAM* in different datasets (TCGA and GEO) had different effects on the survival rates of LUAD patients. Therefore, more experimental evidence and further analysis are required to examine the *ESAM* gene as a potential therapeutic target for LUAD.

The molecular functions of the *NES* gene are protein binding and CCR5 chemokine receptor binding (37, 38). In previous reports, *NES* played an important role in the invasion, metastasis, and prognosis of a variety of tumors, such as glioblastoma, pancreatic cancer, breast cancer, prostate cancer, gastric adenocarcinoma, etc. (39). In addition, Jakub Neradil et al. suggested that *NES* is one of the identifying markers of cancer stem cells (40). In this study, we identified that the *NES* gene was a risk factor in LUAD, and the survival rate of the low-expression group was higher than that of the high-expression group. Additionally, the expression of *NES* for tumor samples was significantly lower than that for normal samples.

Moreover, we analyzed the expression of the prognostic genes in 22 tumors and found that the expression of prognostic genes in different tumors is not completely consistent. For LUAD and LUSC, the expression of the *NES* gene and *ESAM* gene of tumor samples was lower than that of normal samples. In addition to NSCLC, the *NES* gene was also less expressed in tumor samples of KICH, KIRP, PAAD, BLCA, PRAD, KIRC, BRCA, and UCEC. The *ESAM* gene was also less expressed in tumor samples of KICH, KIRP, CESE, THCA, BRCA and UCEC.

Therefore, we have provided a reliable analysis method for screening potential biomarkers of disease. We found that two new biomarker genes play an important role for pan-cancer, and the *NES* gene may be a potential target gene for diagnosis or treatment of LUAD.

## Conclusions

In this study, we used three types of omics data to predict prognostic genes for LUAD and LUSC. Based on the analysis methods in this paper, ten prognostic genes were identified and eight of them have been proven. This implies that our method was feasible. The two novel biomarker genes were gene expression down-regulation and DNA methylation up-regulation and were simultaneously regulated by *miR-122* and *miR-154*. The two biomarker genes of tumor samples were lowly expressed in tumor samples of LUAD, LUSC, KICH, KIRP, BRCA, and UCEC. Unfortunately, we did not find significant differences in expression levels among different stages of NSCLC development.

In summary, we have provided a reliable analysis method for predicting biomarker genes of complex diseases. The two novel biomarker genes play an important role in pan-cancer, and the *NES* gene can be used as a potential target gene for future detection, treatment, and prognosis research of NSCLC.

## Materials and Methods

### Data Acquisition

The RNA-Seq datasets and microRNA-Seq datasets were downloaded from the TCGA database (https://portal.gdc.cancer.gov/). For the lung adenocarcinoma (LUAD) dataset, the RNA-Seq data included 484 lung adenocarcinoma samples and 56 paraneoplastic samples, and the microRNA-Seq data included 485 lung adenocarcinoma samples and 44 paraneoplastic samples. For the lung squamous cell carcinoma (LUSC) dataset, the RNA-Seq data included 474 lung squamous cell carcinoma samples and 48 paraneoplastic samples, and the microRNA-Seq data included 455 lung squamous cell carcinoma samples and 44 paraneoplastic samples. The values of the RNA-seq dataset included COUNT and FPKM; COUNT values were used to obtain differential expression genes, and FPKM values were used for further analyses.

The DNA methylation signal value matrixes were downloaded from the UCSC database (https://xena.ucsc.edu/) (41), which includes 448 LUAD samples, 32 paraneoplastic samples, 346 LUSC samples, and 39 paraneoplastic samples.

The single nucleotide polymorphisms (SNPs) datasets were downloaded from TCGA database. The SNP-related data included 481 and 431 samples of LUAD and LUSC, respectively.

GSE50081 (42), GSE29013 (43), and GSE37745 (44) were downloaded from the GEO database (https://www.ncbi.nlm.nih.gov/gds) through the GEOquery package (45). All of the datasets are from *Homo sapiens*, and the data platform is GPL570. Among the datasets, GSE50081 contains 127 LUAD samples and 42 LUSC samples; GSE29013 contains 30 LUAD samples and 25 LUSC samples; and GSE37745 contains 106 LUAD samples and 66 LUSC samples. We combined and analyzed three datasets related to NSCLC in the GEO database and obtained a total of 263 LUAD samples and 133 LUSC samples. The merged data were standardized using the Limma package (46), and the differences between samples before and after standardization were compared through boxplots (Figure S6).

### Identification of Differentially Expressed Information

The RNA-Seq data and microRNA-Seq data were normalized by a negative binomial-distribution-based approach (47), presented by PCA (Figure S7). For DNA methylation signal value matrix data, the probes (signal values) were filtered when they met the following conditions: detection of failed probes (*P* > 0.01), probes counting less than 3 in at least 5% of samples, probes at non-GpC sites, SNP-associated probes (48), mapping to multiple location probes (49), and sex chromosome probes (50). A cluster analysis was performed on the filtered samples (Figure S8).

The filtering and clustering of the matrix data of DNA methylation signal values were both performed using the ChAMP package (51).

The RNA-Seq data were analyzed using EdgeR (52), and the thresholds for screening differentially expressed genes were *P*-value < 0.05 and |log_2_FC| > 1. In this way, differential expression coding genes hereinafter referred to as differential expression genes (DEGs), and differential long non-coding RNA (DElncRNAs) were screened.

Differential analysis of the DNA methylation signal value matrix was performed using ChAMP. Differential methylation probes (DMPs) with *P*-value < 0.05 and |log_2_FC| > 0.1 were identified. To locate the differential methylation genes (DMGs), the value of log_2_FC in the DMPs was averaged.

The microRNA expression matrix was analyzed using the EdgeR package with *P*-value < 0.05 and |log_2_FC| > 1 to screen for differential microRNA (DEmiRNAs).

### ceRNA Network

The StarBase database (https://starbase.sysu.edu.cn/index.php) (53) provides regulatory relationships for miRNA–lncRNA, miRNA–pseudogene, miRNA–sncRNA, and miRNA–mRNA interactions. We obtained the regulatory relationships between miRNA–mRNA and miRNA– lncRNA in this database as a way to predict the regulatory relationships of DEGs, DElncRNAs, and DEmiRNAs in the ceRNA network and visualized the regulatory relationships using Cytoscape software (54).

### GO and KEGG Enrichment Analysis

To investigate the enrichment of potential prognostic genes in gene functions and pathways, an enrichment analysis was performed. A critical value of FDR < 0.05 was considered statistically significant in the gene ontology (GO) annotation analysis and Kyoto Encyclopedia of Genes and Genomes (KEGG) pathway enrichment analysis of potential prognostic genes using the clusterProfiler package (55).

### Weighted Gene Correlation Network Analysis

The WGCNA package was used to identify co-expressed gene modules and explore the relationship between gene networks and phenotypes (56). The minimum module gene number was set to 50, and the optimal soft threshold was 5 (LUSC) or 6 (LUAD). We were able to identify the co-expression modules of potential prognostic genes in different datasets.

### Protein-Protein Interactions Network Identifies Hub Genes

The STRING (Search Tool for Retrieval of Interacting Genes) online tool (57) was used to predict protein–protein interactions (PPIs), and these were visualized using Cytoscape software (version 3.8.2). In addition, the Molecular Complex Detection (MCODE) plugin in Cytoscape software was applied to explore the hub genes (degree cutoff = 2, k-core = 2, and node score cutoff = 0.2) (58).

### LASSO Regression Model to Construct Diagnostic Model and Screen for Disease Signature Genes

To construct a diagnostic model of NSCLC and screen out disease-characteristic genes, the least absolute shrinkage and selection operator (LASSO) regression model was constructed for hub-genes using the glmnet package (59). In the process of model construction, the characteristics were screened, and the best model was selected to construct the NSCLC risk model. Finally, the genes in the risk model were determined to be NSCLC prognostic genes. In the present study, LUAD and LUSC data were randomly divided into two groups, the training group (80%) and the test group (20%), using the caret package. The accuracy of the diagnostic model was determined by the timeROC package (60).

The tumor samples were also divided into high and low groups according to the optimal cutoff values of risk score, and the optimal cutoff values were calculated using the surv_cutpoint function of survival package.

### Statistical Analysis

R (Version 4.1.1) was used for the statistical analysis and some of the plots in this paper. The *Kolmogorov–Smirnov* test and *chi-squared* test were used to compare the types of variation in the differential gene with other genes. For the comparison of two groups of continuous variables, the statistical significance of normally distributed variables was estimated by an independent *t*-test. The differences between non-normally distributed variables were analyzed using the *Wilcoxon rank sum* test. All statistical *P* values were two-sided and statistically significant at a *P*-value < 0.05. The Kaplan–Meier survival curve was drawn to demonstrate the survival analysis results; genes were divided into a high-expression group and a low-expression group by the optimal cutoff values of expression. Univariate Cox analysis was performed for genes with *P* < 0.05 in the survival analysis. The analysis results are presented in the forest map.

## Abbreviations

NSCLC: non-small cell lung cancer
LUSC: lung squamous cell carcinoma
LUAD: lung adenocarcinoma
BLCA: bladder urothelial carcinoma
BRCA: breast invasive car-cinoma
CESC: cervical squamous cell carcinoma and endocervical adenocarcinoma
CHOL: cholangiocarcinoma
COAD: colon adenocarcinoma
ESCA: esophageal carci-noma
HNSC: head and neck squamous cell carcinoma
KICH: kidney chromophobe
KIRC: kidney renal clear cell carcinoma
KIRP: kidney renal papillary cell carcinoma
LIHC: liver hepatocellular carcinoma
PAAD: pancreatic adenocarcinoma
PCPG: phe-ochromocytoma and paraganglioma
PRAD: prostate adenocarcinoma
READ: rectum adenocarcinoma
SARC: sarcoma
STAD: stomach adenocarcinoma
THCA: thyroid carcinoma
THYM: thymoma
UCEC: uterine corpus endometrial carcinoma
DEGs: differentially expressed genes
DMGs: differentially methylated genes
DMPs: differ-ential methylation probes
DElncRNAs: differential long non-coding RNAs
DEmiRNAs: differential microRNAs.

## Declarations

### Ethics approval and consent to participate

Not applicable.

### Consent for publication

Not applicable.

### Availability of data and materials

All data generated or analyzed during this study are included in this published article and its supplementary information files.

### Authors’ contributions

MYM and KXS conceived the study. JX and HBH performed the data analysis. MYM, JX, and HBH wrote the manuscript. All authors read, improved, and approved the final manuscript.

## Acknowledgments

Not applicable.

